## Supplemental_Information for "Fast and versatile sequence-independent protein docking for nanomaterials design using RPXDock"

#### Bodies

The body class uses pyrosetta to access the pose and initial coordinates of the input .pdb files for a particular docking trajectory. From the pyrosetta pose, the body class stores chain, sequence, secondary structure, and backbone positional information of the asymmetric unit. In this class, any user inputs to allow only certain portions of the pose for docking are also stored (`--allowed_residues`, `--term_access`. Only the initial coordinates of the pyrosetta pose are stored in the body class, while the transformation matrix generated by the search is applied to the initial coordinates of the starting pose. Backbone positional information derived from the transforms are stored as clouds of points during the hierarchical search. The body class checks for clashes between transformed backbones by looking for intersections between these clouds of points at each level of the bounding volume (`intersect_range`, `intersect`, and `clash_ok`). At lower resolutions, the clouds of points are smoother and larger, and at higher resolutions, the clouds of points are smaller. Pairs of contacting positions and secondary structure elements (`contact_pairs`) and counts of contacting pairs (`contact_count`) are also evaluated in the body class.

#### Search

The search module contains the core code controlling the search process. It contains the two fundamental search methods, hierarchical and grid, the geometric specifications (*spec*) for each architecture, the allowed degrees of freedom (*sampler*), and a module for each docking application depending on the architecture and number of bodies. Which module is used depends on the specific architecture and is called by `dock.py`. They are *asym*, *cyclic*, *onecomp*, and *multicomp*. There are also the special-use search applications for one-dimensional (*helix*), and stacking (*axle*) architectures. Finally, the *search* module contains a *result* object which defines the *result* class and associated functions.

Each module type (*asym*, *cyclic*, *onecomp*, *multicomp,* etc.) has a *make function*, e.g., `make_multicomp()` and one or more *evaluator functions*. The *make function* takes as required arguments a body or bodies, a spec, a motif-score hash-table (hscore), a search method (default `hier_search`), and a sampler (default None). The sampler is `hier_multi_axis_sampler()` for multicomp and `hier_axis_sampler()` for onecomp. *Make functions* manage the execution of the docking process by setting up the evaluator function, which performs a redundancy check and calls any specified filters. The evaluator function, in conjunction with the search method, will evaluate the transforms and scores from the sampler at each search resolution and return the top-scoring transforms to be expanded in the next level of search resolution, affecting the docking trajectory. Finally, at the end of docking, the *evaluator function* generates and returns a result object. Result objects are described in further detail in the main text.

#### Score

The `sasa_priority` score function takes into consideration both the quality of the motifs in the interface, and also how far the interface is from a desired size. To develop this function, we generated a predictive model by docking a set of oligomeric scaffolds in all two-component polyhedral group architectures using the `stnd` score function and designed the novel protein-protein interfaces of a random selection of docks using the Rosetta software suite to obtain buried SASA scores (Bale et al. 2016). The resulting model fit to the relationship is

$SASA = 29.1 * ncontact + 282, R^2 = 0.634$. Because `ncontact` correlates strongly with the computationally measured interface size, `SASA`, we parameterized an `ncontact` score term with respect to `SASA` over a range of plausible interface sizes and standard deviations (**Fig. S2**).

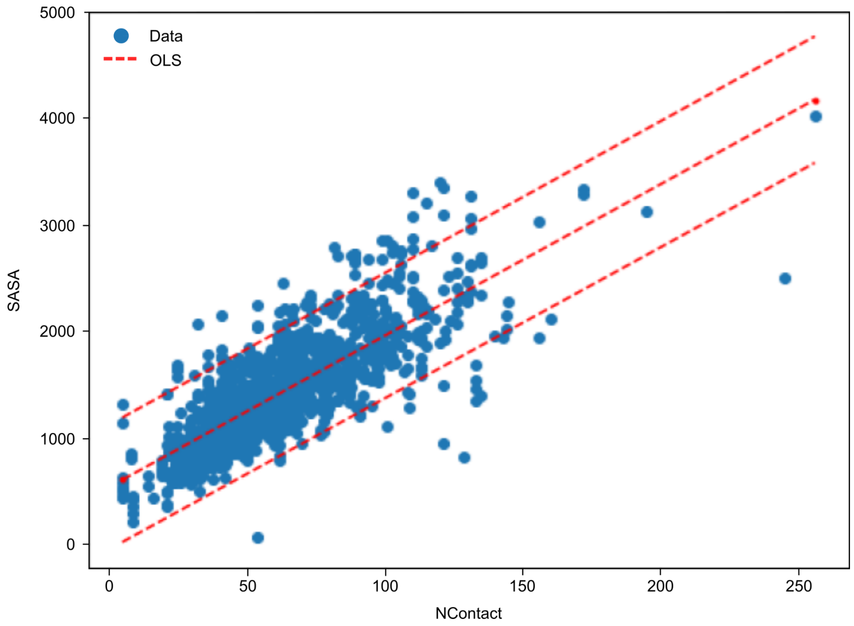


**Figure S2: NContact and SASA are highly correlated.** As such, we parameterized an ncontact score term with respect to computationally measured interface size, SASA.

We fit the correlation between the distribution mean and mode of computationally predicted SASA with a linear regression, and the relationship between the standard deviation and the slope of the correlation between the mean and mode of the distribution with a Gaussian decay function. The resulting log-normal distributions have a maximum score at the input SASA, and the score is invariant with respect to the standard deviation (**Fig. S3**).

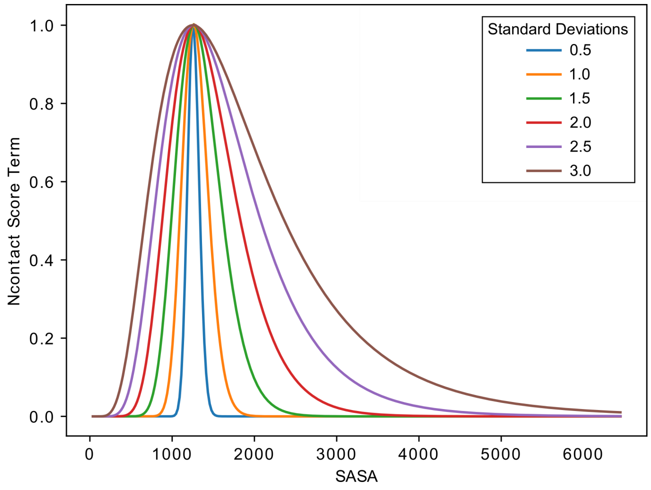


**Figure S3:** Parameterization of an ncontact score term as a function of interface size, SASA, results in a log-normal distribution with a maximum ncontact score term at a user-input SASA regardless of standard deviation.

The final score function contains an *RPX* score term and an ncontact score term: $score\ =\ a * \overline{X}_{RPX}\ +\ b * ln\, N(\mu, \sigma^2)$. The *RPX* score term consists of a scalar multiplier of $\overline{X}_{RPX}$, which refers to the average of the best motifs found across all residue pairs and is used as an approximation of interface quality (Default 1.0). The ncontact score term scores the number of unique contacting pairs (`N`) based on a log-normal distribution set by μ, the desired SASA set by the user, and σ, a user-defined tolerance value that is a scalar multiplier of the standard deviation of the fit error for the correlation between SASA and ncontact, i.e., the accuracy of the prediction for the desired SASA (default 4). Since the *RPX* score term tends to bias towards interfaces that have few but very high quality motif pairs, the weight for the ncontact score term, *b*, needs to be scaled appropriately to overcome the $\overline{X}_{RPX}$ tendency towards small interfaces.

To determine the appropriate default value for *b*, we systematically varied it from 0 to 13 and docked a standard set of scaffolds. The top-, middle-, and bottom-ranked docks were designed using `tools/cage_design.xml`, included in the GitHub repository. Increasing the value of *b* made the interface-size bias to the total score more pronounced, but somewhat surprisingly decreased the maximum interface size observed in all docks (**Fig. S4A**). The weighting also had an unexpected effect on the average RPX score: The $\overline{X}_{RPX}$ also decreased around the desired SASA as *b* increased (**Fig. S4B**). Because $\overline{X}_{RPX}$ is calculated using the mean() gather function, as opposed to a sum() as in the `stnd` score function, this result can be interpreted as the ncontact score term weight having a negative impact on the interface quality for a given interface size. This effect converges above an ncontact weighting of 5, although convergence depends on user-defined interface size (**Fig. S4C**).

We also evaluated the 960 top-, middle-, and bottom-scoring docks for each weighting of the ncontact score term and a target SASA of 1125 Å$^2$ against Rosetta design computational filters including ddG < -20 and SASA between 850 Å$^2$ and 1200 Å$^2$ (included in `cage_design.xml`). Despite the apparent decrease in interface quality as a function of increasing ncontact weight, weights of 5 and above resulted in a higher percentage of top-scoring docks passing Rosetta design filters (**Fig. S4D**). There was also almost no difference in computationally estimated ddG or SASA for top docks above an ncontact weight of 3 (**Fig. S4E-F**). In fact, no statistically significant difference between weights could be detected for any computational design metric. Qualitatively the designs from each weighting look similar, with the top dock for each weight converging after weight = 5 (**Fig. S4G**). Therefore, we selected a default ncontact weight of 5 as the most conservative weighting that also maximizes the design success rate.

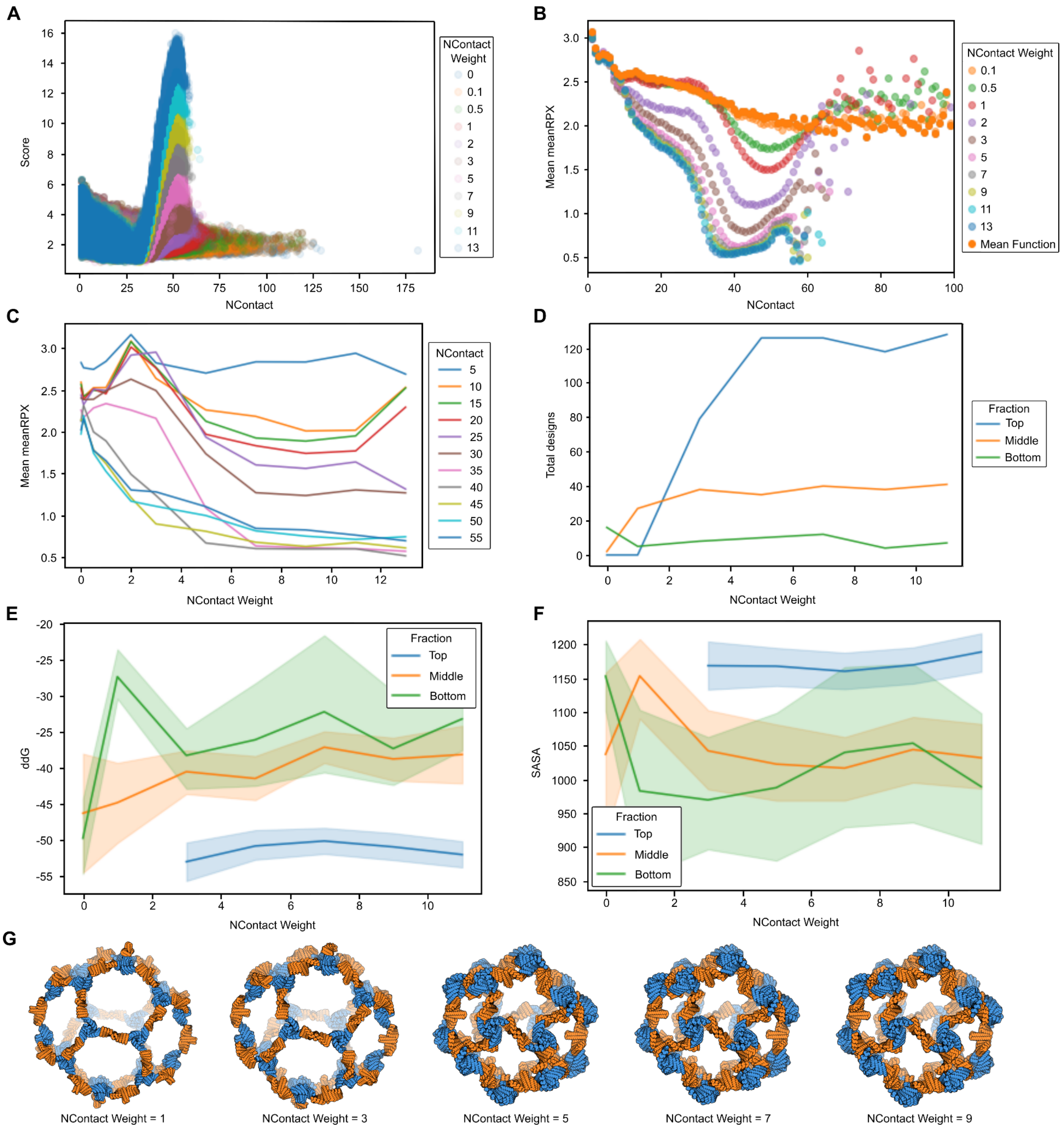


**Figure S4: Empirical derivation of the ncontact score term weight. A.** Score as a function of ncontact across various ncontact weights. **B.** Mean *RPX* as a function of ncontact. **C.** Mean *RPX* as a function of ncontact weighting plotted for interface sizes from Number of unique contacts = 5-55. **D.** Total number of passing designs out of 960 docks for each weighting and fraction. **E-F.** Computational design metrics as a function of ncontact weight for top-, middle-, and bottom-ranked designs for **E.** ddG, and **F.** SASA. **G.** The top dock with I32 icosahedral symmetry for, left to right, ncontact weight 1, 3, 5, 7, 9.

### Filters

The filter module serves two purposes. The first is to filter redundant docks during the search process, described in the Clustering section. The second is to use the `filter()` function to execute an arbitrary number of filters, defined by the user in a filter config file (in .yaml format). This function takes in the body object and transforms, and parses the config file set with the `--filter_config` argument, calling any filters defined in the config file. For all filters, the `filter()` function returns an array of indices for docks passing all filters if the "confidence" configuration is set to *True*. The function also returns any extra data provided by the filters. Available filters are `filter_sasa()` and `filter_sscount()`.

**The SASA filter** attempts to estimate the Solvent Accessible Surface Area buried by the formation of a protein-protein interface. The estimate is based on a linear fit of SASA, calculated by the SASA filter in Rosetta, as a function of the number of unique residues in a docked interface. The `filter_sasa()` function takes as arguments transforms and bodies, as well as parsed keyword arguments from the config file. A full list of options for the SASA filter can be found in **Table S2**.

**The sscount filter** attempts to estimate the number of secondary elements in contact at a protein-protein interface. The filter uses the `secondary_structure_map` class, which maps secondary structure elements (either Helix, Sheet, or Loop), based on user-definitions of each secondary structure element, onto the body object. Secondary structure elements are recorded for a particular body object if a consecutive stretch of identified secondary structure types exceed a given minimum residue length controlled by `min_helix_length`, `min_sheet_length`, and `min_loop_length`. Given the number of unique pairs of residue contacts at a protein-protein interface, the `filter_sscount()`function estimates the number of each secondary structure element type that is in contact at the interface. A full list of options for the sscount filter can be found in **Table S3**.

#### Table S1: All RPXDock command line options

| Option | Default | Description |
|---|---|---|
| `-h, --help` | | Show a full list of RPXDock options |
| **BVH and Bodies** | | |
| `--architecture` | None | architecture to be produced by docking. Can be Cx for cyclic, Dx_y for dihedral, where x is the dihedral symmetry and y is the symmetry of the scaffold, y=2 or y=x, or polyhedral group where the larger axis of symmetry is listed first. Axle docking requires AXLE_X where X is the symmetry of the two input scaffolds. For symmetric axle docking of two scaffolds of different symmetry, use AXLE_1_X_Y where X and Y are the cyclic symmetries of each scaffold. (e.g. AXLE_1_2_3 would dock a dimer against a trimer). Px_yz for layer architectures where Px describes the lattice symmetry and y and z are the scaffold cyclic symmetries. |
| `--inputs1` | None | input structures for single component protocols or first component for 2+ protocols. Can be inputted as a string or list of strings |
| `--inputs2` | None | input structures for the second component for 2+ component protocols. |
| `--inputs3` | None | input structures for the third component for 3+ component protocols. |
| `--max_longaxis_dot_z` | 1.000001 | maximum dot product of the longest input axis (as determined by PCA) with the main symmetry axis (the cosine of the angle between the two axes). Can be used to force cyclic docks to lay flat. |
| `--max_delta_h` | 9999 | maximum difference between cartesian component offsets for multicomponent symmetry axis aligned docking such as cages and layers. |
| **Defining the Search Space** | | |
| `--cart_bounds` | None | cartesian bounds for various protocols. no default as it's protocol specific |

| | | |
|---|---|---|
| `--flip_components` | *True* | list of boolean value(s) specifying if and which components should be allowed to flip in axis aligned docking protocols. |
| `--fixed_rot` | None | list of components (0,1,2,etc) which should be fixed from rotating in hierarchical docking |
| `--fixed_trans` | None | list of components (0,1,2,etc) which should be fixed from translating in hierarchical docking |
| `--fixed_components` | None | list of components (0,1,2,etc) which should be fixed from rotating *and* translating in hierarchical docking |
| `--fixed_wiggle` | None | Similar to fixed_components (input as list 0,1,2,etc) but allows user-inputted translation and rotation wiggling about orientation axis in hierarchical docking |
| `--fw_cartlb` | -5 | Lower bound for fixed_wiggle translation (in Angstroms) |
| `--fw_cartub` | 5 | Upper bound for fixed_wiggle translation (in Angstroms) |
| `--fw_rotlb` | -5 | Lower bound for fixed_wiggle rotation (in degrees) |
| `--fw_rotub` | 5 | Upper bound for fixed_wiggle rotation (in degrees) |
| `--termini_dir1` | None | Restrict sampling for termini orientation of inputs1 either in (*True*) or out (*False*) for each amino or carboxyl termini specified |
| `--termini_dir2` | None | Restrict sampling for termini orientation of inputs2 either in (*True*) or out (*False*) for each amino or carboxyl termini specified |
| `--termini_dir3` | None | Restrict sampling for termini orientation of inputs3 either in (*True*) or out (*False*) for each amino or carboxyl termini specified |
| `--term_access1` | None | Sample docks that pass termini accessibility compliance for inputs1 at either the amino or carboxyl termini, respectively. |
| `--term_access2` | None | Sample docks that pass termini accessibility compliance for inputs2 at either the amino or carboxyl termini, respectively. |
| `--term_access3` | None | Sample docks that pass termini accessibility compliance for inputs3 at either the amino or carboxyl termini, respectively. |
| **Sampling the Search Space** | | |

| | | |
|---|---|---|
| --docking_method | hier | search method to use in docking. available methods may include "hier" for hierarchical search (probably best), “grid” for a flat grid search, and "slide" for a lower dimension grid search using slide moves. Not all options available for all protocols (grid is not available for multicomp docking). |
| --nresl | None (All stages) | number of hierarchical stages to do for hierarchical searches. probably use only for debugging |
| --cart_resl | 10 | resolution of top level cartesian, sometimes ignored, and resolution is taken from hscore data instead |
| --ori_resl | -30 | resolution of top level orientation, sometimes ignored, and resolution is taken from hscore data instead |
| --grid_resolution_cart_angstroms | 1 | cartesian resolution in Angstroms during grid search |
| --grid_resolution_ori_degrees | 1 | rotation orientation resolution in degrees during grid search |
| --beam_size | 100000 | Maximum number of samples for each stage of a hierarchical search protocol (except the first, coarsest stage, which must sample all available positions). This is the most important parameter for determining runtime (aside from number of allowed residues list) |
| **Scoring** | | |
| --function | stnd | score function to use for scoring. Default is stnd score function. Example: stnd, sasa_priority, mean, exp, median. Full list is defined in score/scorefunctions.py |
| --weight_rpx | 1 | score weight of the main *RPX* score component. |
| --weight_ncontact | 0.01 | score weight of each contact (pair of centroids within --max_pair_dist) |
| --weight_sasa | 1152 | Desired SASA used to weight dock scoring for sasa_priority score function |
| --weight_error | 4 | Standard deviation used to calculate the distribution of SASA weighting for sasa_priority score function |
| --hscore_files | 'ilv_h' | *RPX* score files used in scoring for most protocols. defaults to pairs involving only ILV and only in helices. Can be only a path-suffix, |

| | | |
|---|---|---|
| | | which will be appended to --hscore_data_dir. Can be a list of files. |
| --hscore_data_dir | hscore | default path to search for hcores_files |
| --score_only_ss | EHL | only consider residues of the specified secondary structure type when scoring |
| --score_only_sspair | None | only consider pairs with the specified secondary structure types when scoring. may not work in all protocols |
| --allowed_residues1 | None | allowed residues list for single component protocols or first component of 2+ component protocols or the monomeric plug for plug protocol. Takes either nothing (if you leave them out), a single file which applies to all the corresponding inputs, or a list of files which must have the same length as the list of inputs. The files themselves must contain a whitespace separated list of either numbers or ranges. |
| --allowed_residues2 | None | allowed residues list for the second component for 2+ component protocols or the hole for the plug protocol. |
| --allowed_residues3 | None | allowed residues for the third component for 3+ component protocols. |
| --max_pair_dist | 8 | maximum distance between centroids for a pair of residues to be considered interacting. In hierarchical protocols, coarser stages will add appropriate amounts to this distance |
| --ignored_aas | CGP | Amino acids to ignore in scoring |
| --primary_iface_cut | None | score cut for helix primary interface |
| --score_self | *False* | score each interface separately and dump in output pickle |
| **Filtering and Clustering** | | |
| --clashdis | 3.5 | minimum distance allowed between heavy atoms |
| --max_bb_redundancy | 3 | minimum distance between outputs from a single docking run. is more-or-less a non-aligned backbone RMSD |
| --max_cluster | 0 | maximum number of results to cluster (filter redundancy via max_bb_redundancy) for each dock |

| | | |
|---|---|---|
| `--filter_config` | None | Path to a *.yaml file containing the configurations for filters. NOTE: filters only work for cyclic, onecomp, and multicomp docking (ie. not for stacking or asymmetric docking). |
| **Result** | | |
| `--nout_debug` | 0 | Specify number of pdb outputs for individual protocols to output for each search. This is not the preferred way to get pdb outputs, use --nout_top and --nout_each unless you have a reason not to |
| `--nout_top` | 10 | total number of top scoring output structures across all docks. only happens if --dump_pdbs is also specified |
| `--nout_each` | 1 | number of top scoring output structures for each individual dock. only happens if --dump_pdbs is also specified |
| `--dump_pdbs` | *False* | activate output of *.pdb files. |
| `--output_asym_only` | *False* | dump only asu to *.pdbs. Must be used with `--dump_pdbs` also activated. |
| `--output_closest_subunits` | *False* | for two component stuff, output subunit 2 most contacting subunit 1. Must be used with `--dump_pdbs` also activated. |
| `--suppress_dump_results` | *False* | suppress the output of results files |
| `--iface_summary` | min | method to use for summarizing multiple created interfaces into a single score. For example, a three component cage could have 3 interfaces A/B B/C and C/A, or a monomer-based cage plug will have an oligomer interface and an oligomer / cage interface. default is min. e.g. to take the overall score as the worst of the multiple interfaces |
| `--output_prefix` | rpxdock | output file prefix. will output pickles for a base ResPairScore plus --hierarchy_depth hier XMaps |
| `--dont_store_body_in_results` | *False* | reduce result output size and maybe runtime by not including structure information in results objects. will not be able to rescore or output pdbs from results objects. |
| `--loglevel` | INFO | select log level from CRITICAL, ERROR, WARNING, INFO or DEBUG |
| `--use_orig_coords` | *False* | remember and output the original sidechains |

| | | |
|---|---|---|
| | | from the input structures |
| `--symframe_num_helix_repeats` | 10 | number of helix repeat frames to dump |
| **Compute** | | |
| `--ncpu` | All cores or cores available | number of cpu cores to use. defaults to all cores or cores available according to slurm allocation |
| `--nthread` | Single thread and/or ncpu | number of threads to use in threaded protocols |
| `--nprocess` | --`ncpu` | number of processes to use for multiprocess protocols |
| `--trial_run` | *False* | reduce runtime by using minimal samples, smaller score files, etc. |
| `--debug` | *False* | Enable potentially expensive debugging checks |

### Available filter options

##### Table S2: SASA estimate filter parameters

| Parameter | Required Setting | Default | Description |
|---|---|---|---|
| type | filter_sasa | NA | Must be the required setting exactly. |
| confidence | No | *False* | Should the filter remove failing docks from the result table? |
| min_sasa | Yes | 750 | Min interface size. |
| max_sasa | Yes | 1500 | Max interface size. |
| max_dist: | Yes | 8 | Maximum distance between residue pair centroids to count as part of the interface. |
| apply | No | *True* | Return sasa? |
| ncont | No | *False* | Return unique ncontact (instead of normal ncontact). Note if apply and ncont are both false this filter essentially returns ncontact. |

##### Table S3: SScount filter parameters

| Parameter | Required Setting | Default | Description |
|---|---|---|---|
| type | filter_sscount | NA | Must be the required setting exactly. |
| confidence | No | *False* | Should the filter remove failing docks from the result table? |
| min_helix_length | Yes | 4 | Min resis in helix to count as ss element. |
| min_sheet_length | Yes | 3 | Min resis in sheet to count as ss element. |
| min_loop_length | Yes | 1 | Min resis in loop to count as ss element. |
| max_dist | Yes | 8 | Maximum distance between residues to include in SS count. |
| min_element_resis | Yes | 3 | Min interface resis in ss_element to include in ss count. |
| sstype | Yes | "EH" | Types of secondary structure to include in count. |
| min_ss_count | Yes | 3 | If sscount_confidence set, minimum number of ss elements to pass the filter. |
| strict | No | *False* | Require that both pairs of residues in the interface are in an SS element meeting the set criteria. (This is not recommended for standard docking problems). |
